# Heritability and genetic variance of dementia with Lewy bodies

**DOI:** 10.1101/454249

**Authors:** Rita Guerreiro, Valentina Escott-Price, Dena G. Hernandez, Celia Kun-Rodrigues, Owen A. Ross, Tatiana Orme, Joao Luis Neto, Susana Carmona, Nadia Dehghani, John D. Eicher, Claire Shepherd, Laura Parkkinen, Lee Darwent, Michael G. Heckman, Sonja W. Scholz, Juan C. Troncoso, Olga Pletnikova, Ted Dawson, Liana Rosenthal, Olaf Ansorge, Jordi Clarimon, Alberto Lleo, Estrella Morenas-Rodriguez, Lorraine Clark, Lawrence S Honig, Karen Marder, Afina Lemstra, Ekaterina Rogaeva, Peter St. George-Hyslop, Elisabet Londos, Henrik Zetterberg, Imelda Barber, Anne Braae, Kristelle Brown, Kevin Morgan, Claire Troakes, Safa Al-Sarraj, Tammaryn Lashley, Janice Holton, Yaroslau Compta, Vivianna Van Deerlin, Geidy E Serrano, Thomas G Beach, Suzanne Lesage, Douglas Galasko, Eliezer Masliah, Isabel Santana, Pau Pastor, Monica Diez-Fairen, Miquel Aguilar, Pentti J. Tienari, Liisa Myllykangas, Minna Oinas, Tamas Revesz, Andrew Lees, Brad F Boeve, Ronald C. Petersen, Tanis J Ferman, Neill Graff-Radford, Nigel J. Cairns, John C. Morris, Stuart Pickering-Brown, David Mann, Glenda M. Halliday, John Hardy, John Q. Trojanowski, Dennis W. Dickson, Andrew Singleton, for the International Parkinson’s Disease Genomics Consortium, David J. Stone, Jose Bras

## Abstract

Recent large-scale genetic studies have allowed for the first glimpse of the effects of common genetic variability in dementia with Lewy bodies (DLB), identifying risk variants with appreciable effect sizes. However, it is currently well established that a substantial portion of the genetic heritable component of complex traits is not captured by genome-wide significant SNPs. To overcome this issue, we have estimated the proportion of phenotypic variance explained by genetic variability (SNP heritability) in DLB using a method that is unbiased by allele frequency or linkage disequilibrium properties of the underlying variants. This shows that the heritability of DLB is nearly twice as high as previous estimates based on common variants only (31% vs 59.9%). We also determine the amount of phenotypic variance in DLB that can be explained by recent polygenic risk scores from either Parkinson’s disease (PD) or Alzheimer’s disease (AD), and show that, despite being highly significant, they explain a low amount of variance. Additionally, to identify pleiotropic events that might improve our understanding of the disease, we performed genetic correlation analyses of DLB with over 200 diseases and biomedically relevant traits. Our data shows that DLB has a positive correlation with education phenotypes, which is opposite to what occurs in AD. Overall, our data suggests that novel genetic risk factors for DLB should be identified by larger GWAS and these are likely to be independent from known AD and PD risk variants.

## Introduction

Recent studies have highlighted the role of genetics in the common, but often underappreciated, form of dementia that is dementia with Lewy bodies (DLB). Associations with *GBA*, *APOE* and *SNCA* have all been reproducibly reported by independent groups (1–3), and a recent genome-wide association study (GWAS) identified several risk and candidate variants associated with the disease (4). However, GWAS significant single nucleotide polymorphisms (SNPs) often explain only a small proportion of the total heritability estimated (usually from family-based studies) for a given trait, which results in the ‘missing heritability’ issue (5). One of the possible explanations for this issue is that all common SNPs, regardless of their association p-value, contribute to the heritability of complex traits (6–8). However, given that each individual associated marker explains only a small proportion of the genetic variation with little predictive power, methods have been developed to test disorder prediction by summarizing variation across many loci (regardless of association p-values) into quantitative scores. One such approach is the generation of polygenic risk scores (PRSs). PRSs have been successfully applied to Parkinson’s (PD) (9) and Alzheimer’s diseases (AD) (10) and their usefulness will continue to increase as discovery datasets are augmented.

A separate, but related, concept is that of genetic correlation of traits. Here, what is estimated is the genetic covariance between traits that is tagged by common genome-wide SNPs (11). This allows us to identify pleiotropic effects between traits that might be unrelated by any other measurement. We have performed a preliminary study of genetic correlation between DLB and both PD and AD (12), however performing similar analyses with other (even apparently unrelated) traits might provide novel insights for the underlying pathobiology of disease and perhaps for treatments across diseases.

The phenotypic variance of most complex human traits combines the genetic with the environmental variance (13). While the effects of the environment are difficult to ascertain given their complexity and lack of adequate measurements, we are able to determine the genetic variance more accurately. Classically, genetic variance has been partitioned into sources of variation due to additive, dominance and epistatic effects. Additive genetic variance 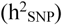 relates to an allele’s independent effect on a phenotype; dominance variance 
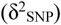 refers to the effect on a phenotype caused by interactions between alternative alleles at a specific locus; epistatic variance refers to the interaction between different alleles in different loci. Most available cohorts for studies of human biology and disease are still underpowered to identify epistatic events, however, additive and dominance variance can be estimated from standard genome-wide genotyping data (14).

Here, using data from the first GWAS in DLB that included haplotype reference consortium (HRC)-imputed genotypes (15), we have estimated the total heritability of this disease. We used a method (GCTA-LDMS) that is unbiased regardless of the minor allele frequency (MAF) and linkage disequilibrium (LD) properties of variants and thus greatly improves on previous estimates (16). Since it has been suggested that heritability estimates may be inflated by non-additive variation (17), we have also estimated the dominance genetic variation in DLB. Additionally, to measure the proportion of variance explained by PRSs from PD and AD in a large DLB cohort, we measured the ability of PRS to discriminate case from control subjects. Lastly, to attempt to derive novel biological insights from unrelated traits, we have performed pairwise genetic correlation analysis of DLB with 235 phenotypes, including cognitive, anthropometric and education traits.

## Results

### Quantifying the genetic heritability of DLB

We applied the GREML-LDMS approach to estimate the proportion of phenotypic variance explained by the HRC-imputed variants for DLB. Results from this approach showed that imputed variants with R^2^ greater than or equal to 0.3 and frequency above 0.1% explained 59.9% (s.e.= 2.1%; p=6.8×10^−6^) of phenotypic variance for DLB. Lower frequency variants explained a large proportion of the phenotypic variance in DLB. This pattern was maintained for the higher quality imputed variants as well (Figure 1, Supplementary Table 1).

**Fig. 1.**
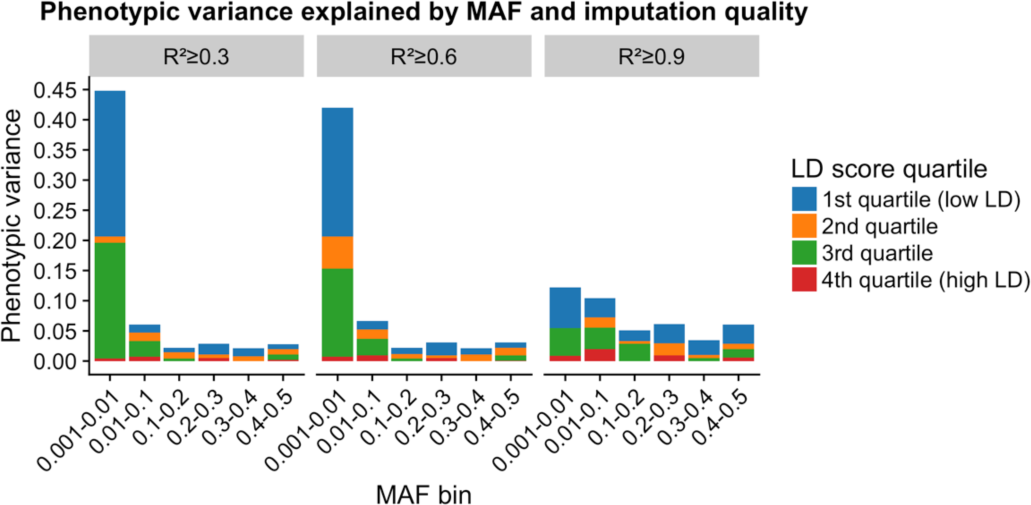
Estimate of the DLB variance explained by HRC-imputed variants by MAF and LD. Segmental LD score increases from the 1st to 4th quartiles. Negative scores are not shown for simplicity but are present in Supplementary Table 1. The estimates of variance explained are from the GREML-LDMS analyses of fitting all the 24 genetic components simultaneously.

To determine if non-additive variance in DLB would explain a subset of the total disease heritability, we calculated the disease dominance variance as implemented in the tool GCTA-GREMLd. This method uses genome-wide data to estimate the additive and dominance genetic relationship matrices (GRMs) and fits both GRMs in a mixed linear model to estimate 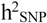 and 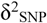 simultaneously. Our results suggest that DLB does not show significant dominance variance with an overall estimate 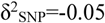 (s.e. = 0.02).

### Polygenic Prediction of Case-Control Status

We applied the PRSs derived from AD and PD data to determine if these would discriminate between DLB and controls. The AD score explained 1.33% of the variance (Nagelkerke’s pseudo-R^2^) and was highly significant (p = 5.8×10^−31^). Performing the same analysis while excluding the *APOE* locus brought the estimate down to 0.14%, while reaching only nominal significance. Using the PD polygenic risk score, we obtained an estimate of 0.37% of the variance in DLB being explained by that score, a result that was also significant (p=6.4×10^−10^). Interestingly, removing the *GBA* locus resulted in only a small reduction in the variance explained by the PD PRS (0.36%; p=1.23×10^−9^) at the best p-value threshold.

The bar plots of DLB variance explained by the AD and PD polygenic risk scores are presented in Figure 2. As expected given these results, DLB cases had on average higher polygenic risk scores than control subjects for both PD and AD (Figure 3).

**Fig. 2.**
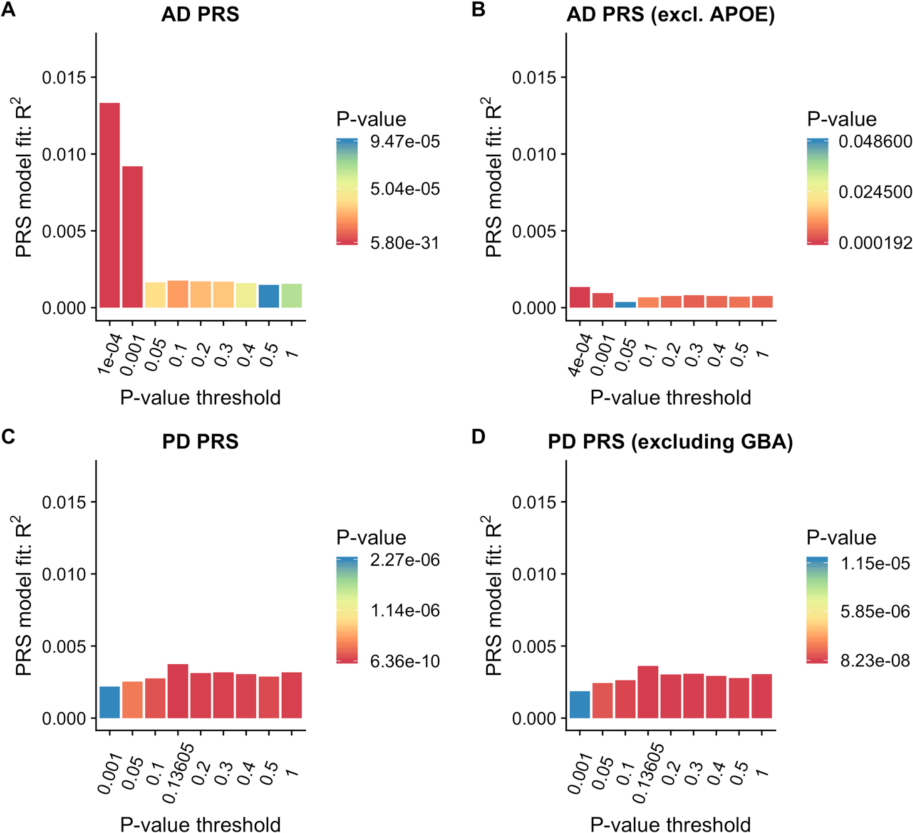
Proportion of variance of DLB case-control status explained by PRSs from AD (A), AD excluding the APOE locus (B), PD (C) and PD excluding the GBA locus (D). The bars represent PRSs calculated for 9 subsets of markers at different p-value thresholds in the original GWAS publications. Best scores for each PRS are presented in (D). R2: Nagelkerke’s pseudo-R2; Threshold: P-value threshold in original GWAS.

**Fig. 3.**
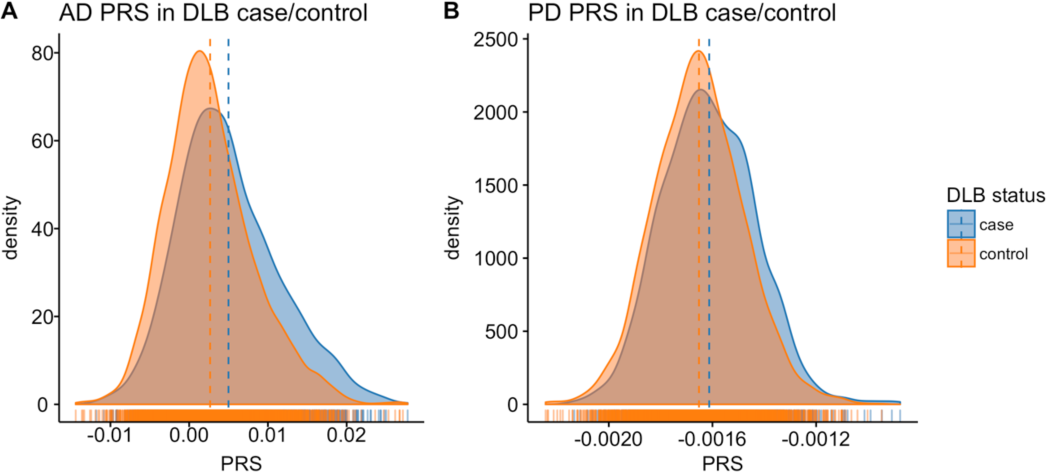
Density distribution of polygenic risk scores (PRS) from AD and PD in DLB case and control subjects. The curves represent the standardized residuals of PRS after adjustment for the first 6 principal components. Blue indicates case subjects; orange indicates case subjects.

### Unbiased genetic correlation

To test whether DLB has a shared genetic etiology with any of 235 other diseases or biomedical relevant traits, we used LD score regression as implemented in LDHub (http://ldsc.broadinstitute.org/ldhub/). This method estimates the degree to which genetic risk factors are shared between pairs of diseases or traits, although it should be noted that it does not inform regarding how this shared genetic etiology arises. We selected the correlations with a p-value <0.01 in DLB and tested these in AD and PD (Figure 4).

**Fig. 4.**
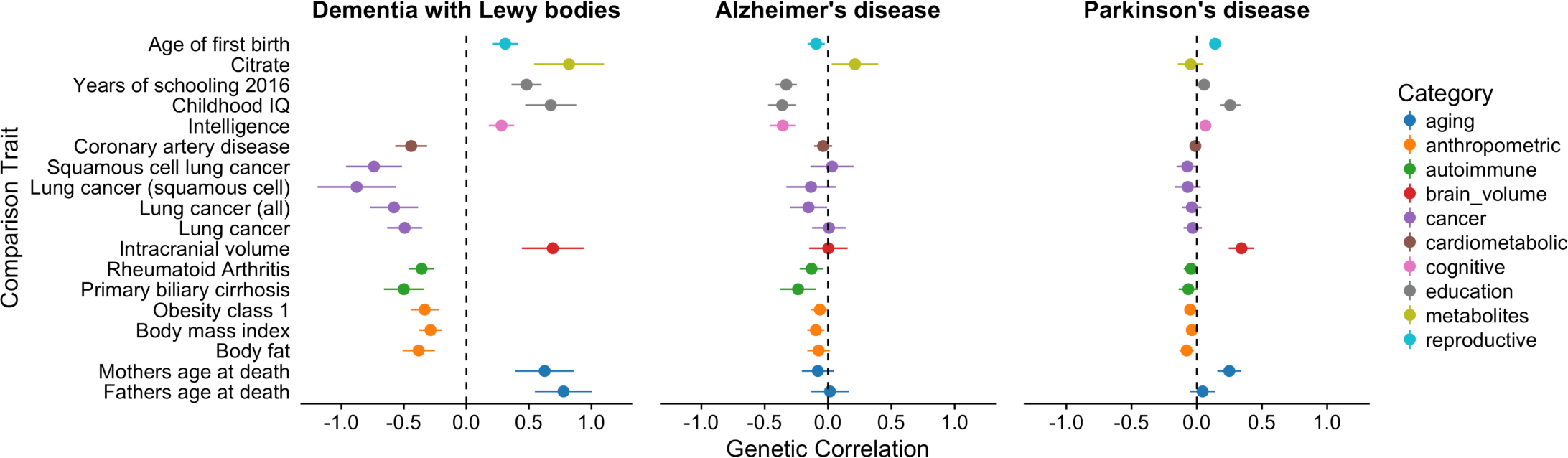
Correlation scores with p-value <0.01 in DLB. Shown are also the scores for those same traits in PD and AD.

The most significant correlation identified between DLB and each of the 235 tested traits was with “Years of schooling” (18) reaching a p-value of 6.32×10^−5^(Bonferroni corrected p-value=0.015) and a correlation estimate (rg) of 0.48 (s.e. = 0.12) (Table 1). Interestingly, these scores were found to be in the opposite direction in AD, but in the same direction in PD (AD: rg=−0.33, p-value=8.87×10^−5^; PD: rg=0.05, p-value=0.07) (Figure 4). A positive correlation was also obtained for “Childhood IQ” (19) in DLB and PD, whereas a negative correlation was identified in AD (DLB: 0.68, p-value=0.0009; AD: rg=−0.36, p-value=0.0011; PD: rg=0.25, p-value=0.0013). Similarly, “Intracranial volume” (20) presented a positive correlation with both DLB and PD, but no discernible correlation with AD (DLB: 0.69, p-value=0.0052; AD: rg=−0.003, p-value=0.96; PD: rg=0.34, p-value=0.0005). Conversely, “Citrate” (21) was positively correlated with both DLB and AD, but had no correlation with PD (DLB: 0.82, p-value=0.0033; AD: rg=−0.21, p-value=0.25; PD: rg=−0.05, p-value=0.63).

**Table 1.**
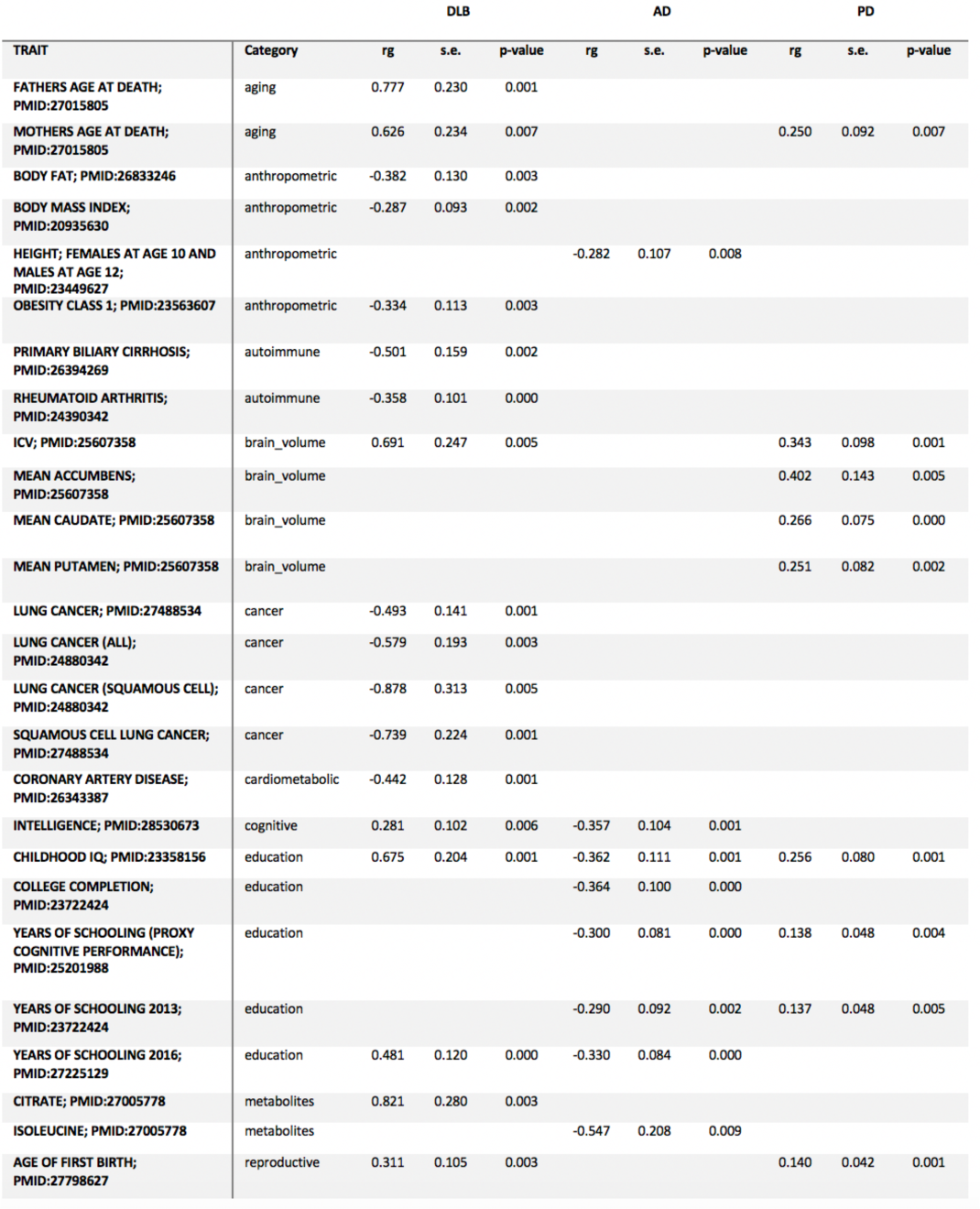
LDHub correlations with p-value <0.01.

## Discussion

With this study we provide more accurate estimates of genetic heritability for DLB, quantify the variance explained by AD and PD polygenic risk and estimate pleiotropy between DLB and over 200 diseases and biomedical relevant traits.

Previous heritability estimates for DLB were calculated based on a smaller cohort genotyped at a relatively smaller number of sites and using GCTA’s GREML-SC (based on a single genetic relationship matrix). These earlier studies provided an estimate of 31% heritability for this disease (12). It is now recognised that GREML-SC may, under certain circumstances (such as causal variants being enriched in regions with higher or lower LD than average or if the causal variants had a different MAF spectrum than the variants sampled), be biased (16). Because of this, we used a recently developed approach that corrects for the LD bias in the estimated SNP-based heritability and that is unbiased regardless of the properties (e.g. MAF and LD) of the underlying causal variants (GCTA GREML-LDMS) (16). We applied this tool to a larger cohort, that was imputed with the most recent imputation panel, providing more detailed genetic information. Using this approach we estimated that all HRC-imputed variants with MAF >0.001 explained 59.9% (s.e= 2.1%) of phenotypic variance for DLB, which is nearly double the previous estimate (12). Our results also show that a large proportion of the variance is explained by variants with lower frequency (MAFs from 0.001 to 0.01). Given that the current version of HRC allows for imputation of variants with frequencies as low as 0.0005 and aggregate R^2^ above 0.5 (15), this indicates that performing GWAS in DLB with increased sample sizes will allow us to identify novel loci involved in conferring risk for disease without the need for large-scale whole-genome sequencing.

One of the explanations for the common issue of “missing heritability” is that non-additive heritability (such as dominance variance or epistatic variance) represents a substantial component of a trait’s total heritable genetic component. Our results suggest that dominance variance has a negligible effect on the genetic heritability of DLB, in line with findings from 79 unrelated traits (14). However, we cannot exclude that epistatic variance plays a role in DLB, given that our cohort is underpowered to detect epistatic events.

Recently, there has been growing interest in the use of PRSs as a way to perform risk prediction in various diseases and these have successfully been applied to AD (10) and PD (9). To determine how much of the phenotypic variance in our DLB cohort can be caused by AD and PD known genetic risk factors, we used PRSs from recent GWAS from each of these diseases. In both cases scores were predictive of case-control status, although explaining only relatively small proportions of variance (0.37-1.33%). In AD, excluding the *APOE* locus greatly reduced the amount of variance explained in DLB (0.14%), which is in accordance with the strong effect that locus has in the risk of both diseases (4, 22). Conversely, excluding the *GBA* locus in PD had only a modest effect, which likely results from the lower frequency in the general population of the variants that comprised this signal compared to *APOE*. Since the amount of variance explained by each of the PRS is relatively small, this adds to the growing body of evidence that suggests that, genetically, DLB is a unique condition and not simply a mix of PD and AD risk factors. These data also confirm the polygenic nature of DLB as well as quantify the amount of variance that polygenic risk from each of those diseases accounts for in DLB.

Given the large number of pleiotropic events that are being identified for a variety of diseases and traits (23, 24), finding correlated conditions opens the door to a better understanding of disease pathobiology and perhaps may even suggest novel therapeutic targets. Assessing the genetic correlation of DLB with over 200 diseases and traits showed correlations that were in the same direction of those seen in PD while others were in the same direction as in AD. It is interesting to note that education scores were positively correlated with DLB, while they have a well established negative correlation with AD (25, 26). Similar positive correlations have been identified for bipolar disorder and autism spectrum disorders (27), as well as for PD in the present data. Also in PD, there is evidence for the presence of increased intracranial volumes when compared to controls (28). Here, supporting those findings, we identify a positive genetic correlation between both PD and DLB with intracranial volume, whereas in AD no evidence for genetic correlation was identified. Interestingly, the anthropometric characteristics obesity, body mass index (BMI) and body fat were negatively correlated with all 3 diseases. For BMI and PD, recent Mendelian randomization results have shown a negative effect (29) which our results replicate and suggest they extend to both AD and DLB. A similar finding was obtained for cancer traits, where lung cancer showed a general negative correlation with the three traits. This agrees with transcriptomic studies that showed that the cancer gene expression profile is almost an opposite mirror image to that of neurodegenerative disease (30). A positive correlation between both DLB and AD with citrate (21) was identified, although this was not the case for PD, where no evidence of correlation was found. Increased plasma levels of citrate have been shown to be associated with increased levels of oxidative stress (31), making it tempting to speculate that in AD and DLB oxidative stress may be involved in the neurodegenerative processes, while in PD it may be more akin to a consequence.

We note several limitations in our study. First, the DLB dataset, despite being the largest to date, is relatively small when compared to other recently published GWAS. This has implications in the statistical power to make novel findings and is reflected in the standard errors of the analyses performed. We are underpowered to detect rare variants and certainly rare variants with small effect sizes. Second, we are unable to provide definitive biological mechanisms underlying the genetic correlations identified. This means that it is possible that for some of the correlations observed, what we are seeing are proxy effects and not direct correlations. Lastly, this study focused on individuals of European/North American descent. It is likely that studies of populations of different ancestries will reveal not only novel loci, but perhaps also novel pleiotropic effects, which could improve our understanding of the pathobiology of DLB.

In summary, we provide updated estimates of the genetic heritability of DLB and show that dominance variance is not a substantial part of the heritability of this disease. We quantify the amount of phenotypic variance in DLB that can be attributed to PD and AD polygenic risk scores and show that this is relatively small. Lastly, we estimate genetic correlations between DLB and over 200 diseases and medically relevant traits, shedding light into the complex relationship between DLB and both PD and AD.

## Materials and Methods

### Sample description

The DLB dataset was previously published (4) and is comprised of 1,216 cases and 3,791 controls, imputed with HRC v1.1 and includes variants with minor allele frequency >= 0.001 and R^2^>=0.3, for a total number of 18.4 million variants (median R^2^=0.92). We used AD summary statistics from the International Genomics of Alzheimer’s Project (IGAP) (22), which is a large two-stage study based upon genome-wide association studies (GWAS) on individuals of European ancestry. In stage 1, IGAP used genotyped and imputed data on 7,055,881 single nucleotide polymorphisms (SNPs) to meta-analyse four previously-published GWAS datasets consisting of 17,008 Alzheimer’s disease cases and 37,154 controls (the European Alzheimer’s disease Initiative – EADI the Alzheimer Disease Genetics Consortium – ADGC, the Cohorts for Heart and Aging Research in Genomic Epidemiology consortium – CHARGE, the Genetic and Environmental Risk in AD consortium – GERAD). PD summary statistics were derived from the International Parkinson’s Disease Genomics Consortium (IPDGC) previously published data and included 13,708 cases and 95,282 controls (32).

### DLB heritability estimates

We used the GCTA-LDMS method to estimate heritability based on imputed data (16, 33) using an imputation quality above 0.3 and a disease prevalence of 0.1%. This method considers the LD-bias that occurs in the SNP-based estimates and is unbiased regardless of the properties of the underlying variants. We calculated segment-based LD scores using a segment length of 200kb (with 100kb overlap between two adjacent segments), which were used to stratify the SNPs into quartiles. We then estimated the genetic relationship matrix (GRM) for each sample using the SNPs in each quartile separately and further stratified by minor allele frequency bins (0.001-0.01, 0.01-0.1, 0.1-0.2, 0.2-0.3, 0.3-0.4, 0.4-0.5). Lastly, we performed restricted maximum likelihood (REML) analysis using the multiple GRMs.

### DLB dominance variance estimates

To estimate the dominance GRM between pairs of individuals, we used genome-wide imputed SNPs as implemented in GCTA-GREMLd (14). This method calculates the additive and dominance GRMs and fits both GRMs in a mixed linear model to estimate additive and dominance variance simultaneously.

### PRS analyses

Determining the polygenic risk of a given phenotype and applying it to another trait is an approach that allows to determine shared genetic aetiology between traits. We calculated PRSs on the base phenotypes (PD and AD), using GWAS summary statistics, and used these as predictors of the target phenotype (DLB) in a regression test. To construct and apply the PRSs we used PRSice v2.1 (34). We performed clumping on the target data by retaining the SNP with the smallest p-value from each LD block (excluding SNPs with r2 > 0.1 in 250kb windows). Each allele was weighted by its effect-size as estimated in the respective study (for PD and AD). Association of PRSs with case-control status was performed with logistic regression, and Nagelkerke’s pseudo-R^2^ was calculated to measure the proportion of variance explained.

### Genetic correlation analysis

To estimate the genetic correlation between DLB and other complex traits and diseases, we used a method based on LD score regression and implemented in the online web utility LDHub v1.9.0 (27, 35). The LD score regression method uses summary statistics from the DLB GWAS and the other available traits, calculates the cross-product of test statistics at each SNP, and then regresses the cross-product on the LD score. After identifying the most significant correlations for DLB (p<0.01), we estimated the correlation of those traits with PD and AD.

## Author Contributions and Notes

RG and JB designed the study and wrote the first draft of the manuscript. JB, RG obtained funding for the study. RG, TO, CKR, JN, JB performed data analyses and interpreted the data. All other authors collected and characterized samples for inclusion in the study. All authors provided critical feedback and helped shape the research, analysis and manuscript. This article contains supporting information online.

## Acknowledgments

Jose Bras and Rita Guerreiro’s work is funded by research fellowships from the Alzheimer’s Society. Tatiana Orme is supported by a scholarship from the Lewy Body Society. This work was supported in part by the National Institutes of Neurological Disease and Stroke. For the neuropathologically confirmed samples from Australia, tissues were received from the Sydney Brain Bank, which is supported by Neuroscience Research Australia and the University of New South Wales, and Dr Halliday is funded by an NHMRC senior principal research fellowship. We would like to thank the South West Dementia Brain Bank (SWDBB) for providing brain tissue for this study. The SWDBB is supported by BRACE (Bristol Research into Alzheimer’s and Care of the Elderly), Brains for Dementia Research and the Medical Research Council. We acknowledge the Oxford Brain Bank, supported by the Medical Research Council (MRC), Brains for Dementia Research (BDR) (Alzheimer Society and Alzheimer Research UK), Autistica UK and the NIHR Oxford Biomedical Research Centre. The brain samples and/or bio samples were obtained from The Netherlands Brain Bank, Netherlands Institute for Neuroscience, Amsterdam (open access: www.brainbank.nl). All Material has been collected from donors for or from whom a written informed consent for a brain autopsy and the use of the material and clinical information for research purposes had been obtained by the NBB. This study was also partially funded by the Wellcome Trust, Medical Research Council (Dr. St. George-Hyslop) and the Canadian Consortium on Neurodegeneration in Aging (E. Rogaeva). Work from Dr. Compta was supported by the CERCA Programme / Generalitat de Catalunya, Barcelona, Catalonia, Spain. The Nottingham Genetics Group is supported by ARUK and The Big Lottery Fund. The effort from Columbia University was supported by the Taub Institute, the Panasci Fund, the Parkinson’s Disease Foundation, and NIH grants NS060113 (Dr Clark), P50AG008702 (P.I. Scott Small), P50NS038370 (P.I.R. Burke), and UL1TR000040 (P.I.H. Ginsberg). Dr Ross is supported by the Michael J. Fox Foundation for Parkinson’s Research, NINDS R01# NS078086. The Mayo Clinic Jacksonville is a Morris K. Udall Parkinson’s Disease Research Center of Excellence (NINDS P50 #NS072187) and is supported by The Little Family Foundation and by the Mangurian Foundation Program for Lewy Body Dementia research and the Alzheimer Disease Research Center (P50 AG016547). The work from the Mayo Clinic Rochester is supported by the National Institute on Aging (P50 AG016574 and U01 AG006786). This work has received support from The Queen Square Brain Bank at the UCL Institute of Neurology; where Dr Lashley is funded by an ARUK senior fellowship. Some of the tissue samples studied were provided by the MRC London Neurodegenerative Diseases Brain Bank and the Brains for Dementia Research project (funded by Alzheimer’s Society and ARUK). This research was supported in part by both the NIHR UCLH Biomedical Research Centre and the Queen Square Dementia Biomedical Research Unit. This work was supported in part by the Intramural Research Program of the National Institute on Aging, National Institutes of Health, Department of Health and Human Services; project AG000951-12. The University of Pennsylvania case collection is funded by the Penn Alzheimer’s Disease Core Center (AG10124) and the Penn Morris K. Udall Parkinson’s Disease Research Center (NS053488). Tissue samples from UCSD are supported by NIH grant AG05131. The authors thank the brain bank GIE NeuroCEB, the French program “Investissements d'avenir” (ANR-10-IAIHU-06). Dr Tienari and Dr Myllykangas are supported by the Helsinki University Central Hospital, the Folkhälsan Research Foundation and the Finnish Academy. This work was in part supported by the Canadian Consortium on Neurodegeneration in Aging (ER). The authors acknowledge the contribution of data from Genetic Architecture of Smoking and Smoking Cessation accessed through dbGAP. Funding support for genotyping, which was performed at the Center for Inherited Disease Research (CIDR), was provided by 1 X01 HG005274-01. CIDR is fully funded through a federal contract from the National Institutes of Health to The Johns Hopkins University, contract number HHSN268200782096C. Assistance with genotype cleaning, as well as with general study coordination, was provided by the Gene Environment Association Studies (GENEVA) Coordinating Center (U01 HG004446). Funding support for collection of datasets and samples was provided by the Collaborative Genetic Study of Nicotine Dependence (COGEND; P01 CA089392) and the University of Wisconsin Transdisciplinary Tobacco Use Research Center (P50 DA019706, P50 CA084724). The data used for the analyses described in this paper were obtained from the database of Genotypes and Phenotypes (dbGaP), at http://www.ncbi.nlm.nih.gov/gap. Genotype and phenotype data for the Genetic Analysis of Psoriasis and Psoriatic Arthritis study were provided by Dr. James T. Elder, University of Michigan, with collaborators Dr. Dafna Gladman, University of Toronto and Dr. Proton Rahman, Memorial University of Newfoundland, providing samples. This work was supported in part by the Intramural Research Program of the National Institutes of Health (National Institute of Neurological Disorders and Stroke; project ZIA NS003154). Tissue samples for genotyping were provided by the Johns Hopkins Morris K. Udall Center of Excellence for Parkinson’s Disease Research (NIH P50 NS38377) and the Johns Hopkins Alzheimer Disease Research Center (NIH P50 AG05146). This study was supported by grants from the National Institutes of Health, the Canadian Institute for Health Research, and the Krembil Foundation. Additional support was provided by the Babcock Memorial Trust and by the Barbara and Neal Henschel Charitable Foundation. JTE is supported by the Ann Arbor Veterans Affairs Hospital.

IGAP: We thank the International Genomics of Alzheimer’s Project (IGAP) for providing summary results data for these analyses. The investigators within IGAP contributed to the design and implementation of IGAP and/or provided data but did not participate in analysis or writing of this report. IGAP was made possible by the generous participation of the control subjects, the patients, and their families. The i–Select chips was funded by the French National Foundation on Alzheimer’s disease and related disorders. EADI was supported by the LABEX (laboratory of excellence program investment for the future) DISTALZ grant, Inserm, Institut Pasteur de Lille, Université de Lille 2 and the Lille University Hospital. GERAD was supported by the Medical Research Council (Grant n° 503480), Alzheimer’s Research UK (Grant n° 503176), the Wellcome Trust (Grant n° 082604/2/07/Z) and German Federal Ministry of Education and Research (BMBF): Competence Network Dementia (CND) grant n° 01GI0102, 01GI0711, 01GI0420. CHARGE was partly supported by the NIH/NIA grant R01 AG033193 and the NIA AG081220 and AGES contract N01–AG–12100, the NHLBI grant R01 HL105756, the Icelandic Heart Association, and the Erasmus Medical Center and Erasmus University. ADGC was supported by the NIH/NIA grants: U01 AG032984, U24 AG021886, U01 AG016976, and the Alzheimer’s Association grant ADGC–10–196728.

IPDGC: We would like to thank all of the subjects who donated their time and biological samples to be a part of this study. This work was supported in part by the Intramural Research Programs of the National Institute of Neurological Disorders and Stroke (NINDS), the National Institute on Aging (NIA), and the National Institute of Environmental Health Sciences both part of the National Institutes of Health, Department of Health and Human Services; project numbers 1ZIA-NS003154, Z01-AG000949-02 and Z01-ES101986. In addition this work was supported by the Department of Defense (award W81XWH-09-2-0128), and The Michael J Fox Foundation for Parkinson’s Research. This work was supported by National Institutes of Health grants R01NS037167, R01CA141668, P50NS071674, American Parkinson Disease Association (APDA); Barnes Jewish Hospital Foundation; Greater St Louis Chapter of the APDA; Hersenstichting Nederland; the Prinses Beatrix Fonds. The KORA (Cooperative Research in the Region of Augsburg) research platform was started and financed by the Forschungszentrum für Umwelt und Gesundheit, which is funded by the German Federal Ministry of Education, Science, Research, and Technology and by the State of Bavaria. This study was also funded by the National Genome Research Network (NGFNplus number 01GS08134, German Federal Ministry for Education and Research), the German Federal Ministry of Education and Research (NGFN 01GR0468, PopGen) and 01EW0908 in the frame of ERA-NET NEURON. In addition, this work was supported by the EU Joint Programme - Neurodegenerative Diseases Research (JPND) project under the aegis of JPND (www.jpnd.eu) through Germany, BMBF, funding code 01ED1406 and iMed - the Helmholtz Initiative on Personalized Medicine. The French GWAS work was supported by the French National Agency of Research (ANR-08-MNP-012). This study was also funded by France-Parkinson Association, the French program “Investissements d’avenir” funding (ANR-10-IAIHU-06) and a grant from Assistance Publique-Hôpitaux de Paris (PHRC, AOR-08010) for the French clinical data. This study was also sponsored by the Landspitali University Hospital Research Fund (grant to SSv); Icelandic Research Council (grant to SSv); and European Community Framework Programme 7, People Programme, and IAPP on novel genetic and phenotypic markers of Parkinson’s disease and Essential Tremor (MarkMD), contract number PIAP-GA-2008-230596 MarkMD (to HP and JHu). The McGill study was funded by the Michael J. Fox Foundation and the Canadian Consortium on Neurodegeneration in Aging (CCNA). This study utilized the high-performance computational capabilities of the Biowulf Linux cluster at the National Institutes of Health, Bethesda, Md. (http://biowulf.nih.gov), and DNA panels, samples, and clinical data from the National Institute of Neurological Disorders and Stroke Human Genetics Resource Center DNA and Cell Line Repository. People who contributed samples are acknowledged in descriptions of every panel on the repository website. We thank the French Parkinson’s Disease Genetics Study Group and the Drug Interaction with genes (DIGPD) study group: Y Agid, M Anheim, A-M Bonnet, M Borg, A Brice, E Broussolle, J-C Corvol, P Damier, A Destée, A Dürr, F Durif, A Elbaz, D Grabil, S Klebe, P. Krack, E Lohmann, L. Lacomblez, M Martinez, V Mesnage, P Pollak, O Rascol, F Tison, C Tranchant, M Vérin, F Viallet, and M Vidailhet. We also thank the members of the French 3C Consortium: A Alpérovitch, C Berr, C Tzourio, and P Amouyel for allowing us to use part of the 3C cohort, and D Zelenika for support in generating the genome-wide molecular data. We thank P Tienari (Molecular Neurology Programme, Biomedicum, University of Helsinki), T Peuralinna (Department of Neurology, Helsinki University Central Hospital), L Myllykangas (Folkhalsan Institute of Genetics and Department of Pathology, University of Helsinki), and R Sulkava (Department of Public Health and General Practice Division of Geriatrics, University of Eastern Finland) for the Finnish controls (Vantaa85+ GWAS data). We used genome-wide association data generated by the Wellcome Trust Case-Control Consortium 2 (WTCCC2) from UK patients with Parkinson’s disease and UK control individuals from the 1958 Birth Cohort and National Blood Service. Genotyping of UK replication cases on ImmunoChip was part of the WTCCC2 project, which was funded by the Wellcome Trust (083948/Z/07/Z). UK population control data was made available through WTCCC1. This study was supported by the Medical Research Council and Wellcome Trust disease centre (grant WT089698/Z/09/Z to NW, JHa, and ASc). As with previous IPDGC efforts, this study makes use of data generated by the Wellcome Trust Case-Control Consortium. A full list of the investigators who contributed to the generation of the data is available from www.wtccc.org.uk. Funding for the project was provided by the Wellcome Trust under award 076113, 085475 and 090355. This study was also supported by Parkinson’s UK (grants 8047 and J-0804) and the Medical Research Council (G0700943 and G1100643). Sequencing and genotyping done in McGill University was supported by grants from the Michael J Fox Foundation and the Canadian Consortium on Neurodegeneration in Aging (CCNA). We thank Jeffrey Barrett and Jason Downing for assistance with the design of the ImmunoChip and NeuroX arrays. DNA extraction work that was done in the UK was undertaken at University College London Hospitals, University College London, who received a proportion of funding from the Department of Health’s National Institute for Health Research Biomedical Research Centres funding. This study was supported in part by the Wellcome Trust/Medical Research Council Joint Call in Neurodegeneration award (WT089698) to the Parkinson’s Disease Consortium (UKPDC), whose members are from the UCL Institute of Neurology, University of Sheffield, and the Medical Research Council Protein Phosphorylation Unit at the University of Dundee. We thank the Quebec Parkinson’s Network (http://rpq-qpn.org) and its members. Mike A. Nalls’ participation is supported by a consulting contract between Data Tecnica International and the National Institute on Aging, NIH, Bethesda, MD, USA, as a possible conflict of interest Dr. Nalls also consults for Illumina Inc, the Michael J. Fox Foundation, University of California Healthcare and Genoom Health among others. This work was supported by the Medical Research Council grant MR/N026004/1.

## References

1. Nalls,M.A., Duran,R., Lopez,G., Kurzawa-Akanbi,M., McKeith,I.G., Chinnery,P.F., Morris,C.M., Theuns,J., Crosiers,D., Cras,P., et al. (2013) A multicenter study of glucocerebrosidase mutations in dementia with Lewy bodies. JAMA Neurol., 70, 727–735.

2. Hardy,J., Crook,R., Prihar,G., Roberts,G., Raghavan,R. and Perry,R. (1994) Senile dementia of the Lewy body type has an apolipoprotein E epsilon 4 allele frequency intermediate between controls and Alzheimer’s disease. Neurosci. Lett., 182, 1–2.

3. Bras,J., Guerreiro,R., Darwent,L., Parkkinen,L., Ansorge,O., Escott-Price,V., Hernandez,D.G., Nalls,M.A., Clark,L.N., Honig,L.S., et al. (2014) Genetic analysis implicates APOE, SNCA and suggests lysosomal dysfunction in the etiology of dementia with Lewy bodies. Hum. Mol. Genet., 23, 6139–6146.

4. Guerreiro,R., Ross,O.A., Kun-Rodrigues,C., Hernandez,D.G., Orme,T., Eicher,J.D., Shepherd,C.E., Parkkinen,L., Darwent,L., Heckman,M.G., et al. (2018) Investigating the genetic architecture of dementia with Lewy bodies: a two‐stage genome-wide association study. Lancet Neurol., 17, 64–74.

5. Manolio,T.A., Collins,F.S., Cox,N.J., Goldstein,D.B., Hindorff,L.A., Hunter,D.J., McCarthy,M.I., Ramos,E.M., Cardon,L.R., Chakravarti,A., et al. (2009) Finding the missing heritability of complex diseases. Nature, 461, 747–753.

6. Yang,J., Lee,T., Kim,J., Cho,M.-C., Han,B.-G., Lee,J.-Y., Lee,H.-J., Cho,S. and Kim,H. (2013) Ubiquitous polygenicity of human complex traits: genome-wide analysis of 49 traits in Koreans. PLoS Genet., 9, e1003355.

7. Lee,S.H., Wray,N.R., Goddard,M.E. and Visscher,P.M. (2011) Estimating missing heritability for disease from genome-wide association studies. Am. J. Hum. Genet., 88, 294–305.

8. Boyle,E.A., Li,Y.I. and Pritchard,J.K. (2017) An Expanded View of Complex Traits: From Polygenic to Omnigenic. Cell, 169, 1177–1186.

9. Escott-Price,V., International Parkinson’s Disease Genomics Consortium, Nalls,M.A., Morris,H.R., Lubbe,S., Brice,A., Gasser,T., Heutink,P., Wood,N.W., Hardy,J., et al. (2015) Polygenic risk of Parkinson disease is correlated with disease age at onset. Ann. Neurol., 77, 582–591.

10. Escott-Price,V., Sims,R., Bannister,C., Harold,D., Vronskaya,M., Majounie,E., Badarinarayan,N., GERAD/PERADES, IGAP consortia, Morgan,K., et al. (2015) Common polygenic variation enhances risk prediction for Alzheimer’s disease. Brain, 138, 3673–3684.

11. Lee,S.H., Yang,J., Goddard,M.E., Visscher,P.M. and Wray,N.R. (2012) Estimation of pleiotropy between complex diseases using single-nucleotide polymorphism-derived genomic relationships and restricted maximum likelihood. Bioinformatics, 28, 2540–2542.

12. Guerreiro,R., Escott-Price,V., Darwent,L., Parkkinen,L., Ansorge,O., Hernandez,D.G., Nalls,M.A., Clark,L., Honig,L., Marder,K., et al. (2016) Genome-wide analysis of genetic correlation in dementia with Lewy bodies, Parkinson’s and Alzheimer’s diseases. Neurobiol. Aging, 38, 214.e7–10.

13. Mackay,T.F. (2001) The genetic architecture of quantitative traits. Annu. Rev. Genet., 35, 303–339.

14. Zhu,Z., Bakshi,A., Vinkhuyzen,A.A.E., Hemani,G., Lee,S.H., Nolte,I.M., van Vliet-Ostaptchouk,J.V., Snieder,H., LifeLines Cohort Study, Esko,T., et al. (2015) Dominance genetic variation contributes little to the missing heritability for human complex traits. Am. J. Hum. Genet., 96, 377–385.

15. McCarthy,S., Das,S., Kretzschmar,W., Delaneau,O., Wood,A.R., Teumer,A., Kang,H.M., Fuchsberger,C., Danecek,P., Sharp,K., et al. (2016) A reference panel of 64,976 haplotypes for genotype imputation. Nat. Genet., 48, 1279–1283.

16. Yang,J., Bakshi,A., Zhu,Z., Hemani,G., Vinkhuyzen,A.A.E., Lee,S.H., Robinson,M.R., Perry,J.R.B., Nolte,I.M., van Vliet-Ostaptchouk,J.V., et al. (2015) Genetic variance estimation with imputed variants finds negligible missing heritability for human height and body mass index. Nat. Genet., 47, 1114–1120.

17. Eichler,E.E., Flint,J., Gibson,G., Kong,A., Leal,S.M., Moore,J.H. and Nadeau,J.H. (2010) Missing heritability and strategies for finding the underlying causes of complex disease. Nat. Rev. Genet., 11, 446–450.

18. Okbay,A., Beauchamp,J.P., Fontana,M.A., Lee,J.J., Pers,T.H., Rietveld,C.A., Turley,P., Chen,G.-B., Emilsson,V., Meddens,S.F.W., et al. (2016) Genome-wide association study identifies 74 loci associated with educational attainment. Nature, 533, 539–542.

19. Benyamin,B., Pourcain,B., Davis,O.S., Davies,G., Hansell,N.K., Brion,M.-J.A., Kirkpatrick,R.M., Cents,R.A.M., Franić,S., Miller,M.B., et al. (2014) Childhood intelligence is heritable, highly polygenic and associated with FNBP1L. Mol. Psychiatry, 19, 253–258.

20. Hibar,D.P., Stein,J.L., Renteria,M.E., Arias-Vasquez,A., Desrivières,S., Jahanshad,N., Toro,R., Wittfeld,K., Abramovic,L., Andersson,M., et al. (2015) Common genetic variants influence human subcortical brain structures. Nature, 520, 224–229.

21. Kettunen,J., Demirkan,A., Würtz,P., Draisma,H.H.M., Haller,T., Rawal,R., Vaarhorst,A., Kangas,A.J., Lyytikäinen,L.-P., Pirinen,M., et al. (2016) Genome-wide study for circulating metabolites identifies 62 loci and reveals novel systemic effects of LPA. Nat. Commun., 7, 11122.

22. Lambert,J.C., Ibrahim-Verbaas,C.A., Harold,D., Naj,A.C., Sims,R., Bellenguez,C., DeStafano,A.L., Bis,J.C., Beecham,G.W., Grenier-Boley,B., et al. (2013) Meta-analysis of 74,046 individuals identifies 11 new susceptibility loci for Alzheimer’s disease. Nat. Genet., 45, 1452–1458.

23. Visscher,P.M., Wray,N.R., Zhang,Q., Sklar,P., McCarthy,M.I., Brown,M.A. and Yang,J. (2017) 10 Years of GWAS Discovery: Biology, Function, and Translation. Am. J. Hum. Genet., 101, 5–22.

24. Guerreiro,R., Brás,J., Hardy,J. and Singleton,A. (2014) Next generation sequencing techniques in neurological diseases: redefining clinical and molecular associations. Hum. Mol. Genet., 23, R47–53.

25. Barnes,D.E. and Yaffe,K. (2011) The projected effect of risk factor reduction on Alzheimer’s disease prevalence. Lancet Neurol., 10, 819–828.

26. Norton,S., Matthews,F.E., Barnes,D.E., Yaffe,K. and Brayne,C. (2014) Potential for primary prevention of Alzheimer’s disease: an analysis of population-based data. Lancet Neurol., 13, 788–794.

27. Bulik-Sullivan,B., Finucane,H.K., Anttila,V., Gusev,A., Day,F.R., Loh,P.-R., ReproGen Consortium, Psychiatric Genomics Consortium, Genetic Consortium for Anorexia Nervosa of the Wellcome Trust Case Control Consortium 3, Duncan,L., et al. (2015) An atlas of genetic correlations across human diseases and traits. Nat. Genet., 47, 1236–1241.

28. Krabbe,K., Karlsborg,M., Hansen,A., Werdelin,L., Mehlsen,J., Larsson,H.B.W. and Paulson,O.B. (2005) Increased intracranial volume in Parkinson’s disease. J. Neurol. Sci., 239, 45–52.

29. Noyce,A.J., Kia,D.A., Hemani,G., Nicolas,A., Price,T.R., De Pablo-Fernandez,E., Haycock,P.C., Lewis,P.A., Foltynie,T., Davey Smith,G., et al. (2017) Estimating the causal influence of body mass index on risk of Parkinson disease: A Mendelian randomisation study. PLoS Med., 14, e1002314.

30. Aramillo Irizar,P., Schäuble,S., Esser,D., Groth,M., Frahm,C., Priebe,S., Baumgart,M., Hartmann,N., Marthandan,S., Menzel,U., et al. (2018) Transcriptomic alterations during ageing reflect the shift from cancer to degenerative diseases in the elderly. Nat. Commun., 9, 327.

31. Convertini,P., Menga,A., Andria,G., Scala,I., Santarsiero,A., Castiglione Morelli,M.A., Iacobazzi,V. and Infantino,V. (2016) The contribution of the citrate pathway to oxidative stress in Down syndrome. Immunology, 149, 423–431.

32. Nalls,M.A., Pankratz,N., Lill,C.M., Do,C.B., Hernandez,D.G., Saad,M., DeStefano,A.L., Kara,E., Bras,J., Sharma,M., et al. (2014) Large-scale meta-analysis of genome-wide association data identifies six new risk loci for Parkinson’s disease. Nat. Genet., 46, 989–993.

33. Yang,J., Lee,S.H., Goddard,M.E. and Visscher,P.M. (2011) GCTA: a tool for genome-wide complex trait analysis. Am. J. Hum. Genet., 88, 76–82.

34. Euesden,J., Lewis,C.M. and O’Reilly,P.F. (2015) PRSice: Polygenic Risk Score software. Bioinformatics, 31, 1466–1468.

35. Zheng,J., Erzurumluoglu,A.M., Elsworth,B.L., Kemp,J.P., Howe,L., Haycock,P.C., Hemani,G., Tansey,K., Laurin,C., Early Genetics and Lifecourse Epidemiology (EAGLE) Eczema Consortium, et al. (2017) LD Hub: a centralized database and web interface to perform LD score regression that maximizes the potential of summary level GWAS data for SNP heritability and genetic correlation analysis. Bioinformatics, 33, 272–279.

